# Decreased synthesis and variable gene transcripts of oxytocin in a domesticated avian species

**DOI:** 10.1101/2021.03.17.435911

**Authors:** Yasuko Tobari, Constantina Theofanopoulou, Chihiro Mori, Yoshimi Sato, Momoka Marutani, Sayaka Fujioka, Norifumi Konno, Kenta Suzuki, Akari Furutani, Shiomi Hakataya, Cheng-Te Yao, En-Yun Yang, Chia-Ren Tsai, Pin-Chi Tang, Chih-Feng Chen, Cedric Boeckx, Erich D. Jarvis, Kazuo Okanoya

## Abstract

The Bengalese finch was domesticated more than 250 years ago from the wild white-rumped munia. Similar to other domesticated species, Bengalese finches show a reduced fear response and have lower corticosterone levels, compared to white-rumped munias. Bengalese finches and munias also have different song types. Since oxytocin (*OT*) has been found to be involved in stress coping and auditory processing, we tested whether the *OT* sequence and brain expression pattern and content differ in wild munias and domesticated Bengalese finches. We identified intra-strain variability in the untranslated regions of the *OT* sequence in Bengalese finches in comparison to the munia *OT*. Several of these changes fall in specific transcription factor binding sites, which show either a conserved or a relaxed evolutionary trend in the avian lineage, and in vertebrates in general. Although *in situ* hybridization in several hypothalamic nuclei did not reveal significant differences in the number of cells expressing *OT* between the two strains, real-time quantitative PCR showed significantly lower *OT* mRNA expression in the diencephalon of the Bengalese finches relative to munias. Our study thus points to a decreased *OT* synthesis in the domestic strain compared with the wild strain in birds. This is an opposite pattern from that found in some domesticated mammals, suggesting that different processes of *OT* function might have occurred in mammals and birds under domestication.

## 1 Introduction

Domesticated animals have been long known to exhibit similarities, such as: loss of pigmentation; rounded faces; smaller teeth and weaker biting force; floppy ears; shorter muzzles; curly tails; smaller cranial capacity and brain size; pedomorphosis; neotenous (juvenile) behavior; reduction of sexual dimorphism (feminization); docility; and reduced aggression and cortisol levels^1–3^. These changes are collectively referred to as “domestication syndrome.” A theoretical analysis by Wilkins, et al. (2014)^3^ proposed that these common changes may be due to reduced numbers and/or delayed migration of neural crest cells during embryogenesis, that end up forming the cells that make up the aforementioned tissues.

White-rumped munia (WRM; *Lonchura striata*) is a songbird species belonging to Passeriformes, which was imported and brought into captivity from China to Japan around 1762 CE^4^. Munias were initially used to foster exotic birds, which requires calmness under captivity and acceptability of non-kin individuals or species^5^. By artificially selecting against aggressive individuals, Bengalese finches (BF; *Lonchura striata* var. *domestica*) evolved as the domesticated strain of munias; they typically have lower levels of the stress hormone corticosterone (CORT), possibly because parenting in small cages requires stress tolerance^6^. The delayed migration of neural crest cells may cause the overall white appearance of the domesticated BFs^6^, relative to the mostly brown plumage of munias that only have a small white patch on the rump. The assumption that the BF is a domesticated strain of the WRM is supported by the fact that F1 hybrids between BFs and WRMs are fertile^4^, their mostly innate and sexually dimorphic distances calls are nearly the same^7^, occasional occurrences of munia-like morphs in BFs^8^, and the avicultural records^5^.

One gene that has been proposed to play a role in domestication is oxytocin (*OT*)^9^. It is produced mainly in the supraoptic nucleus (SO) and the paraventricular nucleus (PVN) of the hypothalamus, from where it is released in the capillaries of the posterior pituitary to then be distributed peripherally, acting as a hormone, or from axon terminals of PVN neurons that innervate many other brain regions containing the *OT* receptor (*OTR*), acting as a neurotransmitter/ neuromodulator^10^. Traditionally, *OT* has been studied in reproductive contexts, including uterine contractions and milk secretion^11^, while more recently it has been shown to be involved in a wider array of functions, including social bonding and stress suppression^12,13^. A potential role in domestication^9,14^ has been based on findings that show different brain *OT* expression patterns in domesticated versus wild mice and rats^15^; strong purifying selection in the *OT* gene in domesticated placental mammals^16^; single nucleotide polymorphisms in *OT* in domestic camelid populations^17^; and differential brain gene expression in the *OTR* in dogs and wolves^18^. Further, *OT* has been shown to be involved in several facets of social cognition^19^, like social communication^20^ and social avoidance^21^, in mammals, through innervation of PVN *OT* neurons in the lateral septum^22,23^ and the nucleus accumbens^24^, among others. Although less extensively studied, *OT*’s role in social behaviors has been also identified in avian species. For example, in zebra finches *OT* has been shown to regulate aggression, affiliation, allopreening and partner preference^25–27^.

We thus hypothesized that *OT* modifications may have played a role in the domestication of BFs. To test this hypothesis, we compared the nucleotide sequence and hypothalamic *OT* expression between BFs and WRMs. We use the term *OT* for birds, as opposed to the commonly used mesotocin^25^, following the proposal for a universal vertebrate nomenclature for *OT*, its sister gene vasotocin (*VT*), and their receptors (*OTR* and *VTRs*), based on vertebrate-wide synteny results^28^. We found specific nucleotide changes between the BF and the munias in the 5’-untranslated region (UTR) and the 3’-UTR of the *OT* mRNA, and lower OT expression in the diencephalon of the BFs.

## 2 Materials and Methods

All procedures were conducted in accordance with: the guidelines and regulations approved by the Ethical committee of Azabu University; the Guidelines for Animal Experiments of Azabu University or of the RIKEN Animal Experiments Committee; the RIKEN BSI guidelines or the guidelines approved by the Animal Experimental Committee at the University of Tokyo, all of which are in accordance with the relevant guidelines and regulations approved by the Animal Care and Use Committee of Endemic Species Research Institute (ESRI).

### 2.1 Animals used for RNA isolation and *in situ* hybridization (ISH)

The origin and history of the animals used in this study are listed in Supplementary Table 1. Adult male BFs (n = 5) were obtained from our breeding colony at the RIKEN-BSI, Japan. These birds were not siblings, half-siblings or progeny. Adult male WRMs (n = 2) were kept for more than 1 year after they had been purchased from pet breeders or bred in our laboratory. The birds were kept in group housing, under a 12-h light/dark cycle (lights on at 8:00 am) and were supplied ad libitum with mixed seeds and water. The birds were caught by hand and then decapitated between 9:00 am-12:00 pm. The brains of the birds were frozen in optimum cutting temperature, embedded in medium (Sakura Finetek, Tokyo, Japan), and frozen on dry ice. Harvested brains were stored at −80°C until RNA extraction for cDNA cloning.

Using mist nets, wild WRMs (n = 2 males) were captured in Huben (23°43’51”N, 120°37’58”E or 23°43’51”N, 120°37’51”E) Yunlin, Taiwan on March 14, 2019. After capture, birds were transported to Endemic Species Research Institute (ESRI) and brought into the aviary. These two birds were killed by rapid decapitation on March 14 or 15, 2019. The wild munia brains were dissected; the hypothalamus was harvested and placed into RNA later (Ambion, Bicester, UK) and stored overnight at 4°C, shipped on dry ice to Azabu University and stored at −80°C until RNA extraction for cDNA cloning.

Additional six BFs and five WRMs were used for ISH. Both BFs and WRMs were born in our laboratory at RIKEN and had been kept together with conspecifics until decapitation. The BF approximate ages were 120-240 pdh. The WRM ages were over 90 pdh. Both BFs and WRMs had reached sexual maturity. All the birds were kept under a 12-h light/dark cycle (lights on at 8:00 am) and were supplied ad libitum with mixed seeds and water. The birds were caught by hand and then decapitated between 9:00 am-12:00 pm. The brains of the birds were frozen in optimum cutting temperature, embedded in medium (Sakura Finetek,) and frozen on dry ice. Harvested brains were stored at −80°C. The whole brains of BFs (n = 6) and WRMs (n = 4) were used for ISH. For one WRM, the lateral two-thirds of the right hemisphere had already been used for another study; thus, the left hemisphere and the medial one-third of the right hemisphere were used for ISH.

### 2.2 Animals used for quantitative PCR (qPCR) and enzyme immunoassay (EIA)

Thirteen wild male WRMs were captured in Huben (23°43’51”N, 120°37’58”E or 23°43’51”N, 120°37’51”E) or Yuchi village (23°53’31”N, 120°53’31”E), in Yunlin, Taiwan on March 12-14, 2019. After capture, birds were transported to ESRI and brought into the aviary and randomly assigned into cages in one room. The birds were kept for at least 1 week after they had been captured from the wild. Because capturing and transporting the birds would impose more stress on wild-captured WRMs, one week of accommodation under laboratory conditions was provided to minimize stress effects on them. The birds were kept in group housing and on the natural photoperiod (Day length was 12 h 23 min; sunrise and sunset occur at 0524 and 1811, respectively) and were supplied ad libitum with mixed seeds and water. They sang actively, indicating that they were not over stressed by this time. In order to reduce stimuli disparities between subjects, the day before they were decapitated, their cages were shaded with cardboard since 6:00 pm. During March 20-22, all birds were killed by rapid decapitation between 8:00-9:00 am. They weighed from 9.3 to 12 g (mean ± SD: 10 ± 0.64 g), and observation of their large and mature testes after decapitation suggested they were sexually mature males. The brains of the birds were frozen in optimum cutting temperature and embedded in medium (Sakura Finetek) on dry ice. Harvested brains were stored at −80°C and shipped on dry ice to Azabu University and stored at −80°C until RNA (n = 6) and peptide (n = 7) extraction. The birds were treated and handled in accordance with all appropriate animal care guidelines and permits (ESRI). Six male BFs were used for qPCR. These birds were obtained from our breeding colony at the University of Tokyo, Japan and were around two years old (phd 506-794). All six birds were placed in one cage. They were kept under a 14-h light/10-h dark cycle (lights on at 8:00 am) and were supplied ad libitum with mixed seeds and water. They also sang actively. In order to keep the same condition as in Taiwan, the day before they were decapitated, their cages had been covered with cardboard since 6:00 pm. They were killed by rapid decapitation between 8:00-9:00 am. The brains were frozen on dry ice and stored at −80°C until RNA extraction. Frozen brains were dissected into the cerebrum, diencephalon, midbrain and cerebellum immediately before RNA extraction. The cerebellum was dissected out. The diencephalon was dissected by two coronal cuts at the level of the tractus septopalliomesencephalicus (rostral edge of the preoptic area) and the oculomotor nerves (caudal edge of hypothalamus), one parasagittal cut placed 2 mm lateral to the midline and one horizontal cut 5 mm above the floor of the brain, including the hypothalamus. The optic tectum was collected as the midbrain. The posterior telencephalon including the septum was collected as the cerebrum.

Eight BFs were prepared for EIA. They were kept for 19 days at our aviary in Azabu University, Japan, after they had been purchased from a pet breeder. They were around two years old and sang actively. Four birds were placed in one cage, kept under a 12-h light/dark cycle (lights on at 6:00 am) and were supplied ad libitum with mixed seeds and water. The day before they were decapitated, their cages had been covered with shaded cloth since 6:00 pm. One bird was omitted because of cataracts. Seven birds were killed by rapid decapitation between 8:00-9:00 am. Brains were dissected by two coronal cuts at the level of the tractus septopalliomesencephalicus (rostral edge of the preoptic area) and the oculomotor nerves (caudal edge of hypothalamus), one parasagittal cut placed 2 mm lateral to the midline and one horizontal cut 7 mm above the floor of the brain, including the hypothalamus and the bed nucleus of the stria terminalis. Harvested brain tissue blocks were stored at −80°C until peptide extraction for EIAs.

### 2.3 Molecular cloning of songbird OT cDNA

Total RNA was extracted from the hypothalamus of the BF and WRM; it was purified using RNeasy Lipid Tissue mini kit (Qiagen, Hilden, Germany). The unknown cDNA sequences, including 3’- and 5’-UTRs of these three songbirds’ OT genes, were analyzed by PCR amplification of 3’- and 5’-RACE fragments. The first-strand cDNA for 3’-RACE was prepared by incubation of total RNA with adaptor-oligo(dT)_18_ primers and SuperScript III reverse transcriptase (Invitrogen, Carlsbad, CA, USA) at 50°C for 50 min. The PCR primers listed in Supplementary Table 2 were designed based on predicted sequences retrieved from the genome assembly (Taeniopygia_guttata-3.2.4, July 2008^28^). The RACE fragments were amplified using two rounds of PCR with gene-specific primers and adaptor primers. PCR amplification of 3’-RACE fragments for OT consisted of initial denaturation at 94°C for 5 min; followed by 25 cycles of 30 s at 94°C, 30 s at 55°C, and 1 min at 72°C; and final elongation at 72°C for 5 min. The reaction was performed in a 25-μl mixture containing 0.3 μg cDNA, 0.2 mM dNTPs, 0.4 μM each forward and reverse primers, and 2.5 U Ex Taq polymerase with its buffer (TaKaRa, Shiga, Japan). The template cDNA was reverse transcribed with 5’-RACE RT primer and SuperScript III reverse transcriptase (Invitrogen), followed by poly(A) tailing of the cDNA with dATP and terminal transferase (Roche Diagnostics, Rotkreuz, Switzerland). PCR amplification of 5’-RACE fragments for OT consisted of initial denaturation at 98°C for 5 min; followed by 30 cycles of 30 s at 98°C, 30 s at 55°C, and 1 min at 72°C; and a final elongation at 72°C for 5 min. The reaction was performed in a 25-μl mixture comprising 150 ng cDNA, 0.4 mM dNTPs, 0.4 μM each forward and reverse primers, and 2.5 U Ex Taq polymerase with its buffer.

To confirm the ORF sequence, the PCR reaction was performed in a 25-μl mixture containing 0.4 mM dNTPs, 0.4 μM each forward and reverse primers, and 2.5 U Ex Taq polymerase with its buffer, as well as 250 ng cDNA that had been reverse transcribed with SuperScript IV VILO Master Mix (Invitrogen). PCR reaction conditions were as follows: 98°C for 1 min; 30 cycles of 98°C for 10 s, 55°C for 30 s, 72°C for 1 min; and 72°C for 5 min.

PCR products were subcloned into pGEM-T easy vectors (Promega, Madison, WI, USA). The resultant plasmids were sequenced commercially (Fasmac, Atsugi, Japan). The ORF sequences obtained were submitted to DDBJ/EMBL/GenBank with accession numbers LC489419 and LC489420. We additionally used the genomic sequences of the OT we found in the publicly available scaffold-level and chromosome-level BF genome assemblies (LonStrDom1; RefSeq assembly accession: GCF_002197715.1; LonStrDom2, GCF_005870125.1) (Supp. Data S1 and S2).

### 2.4 Database Analysis

A putative signal peptide was predicted by using SignalP 3.0 (http://www.cbs.dtu.dk/services/SignalP/). The chromosomal location and strand orientation of identified BF genes were determined using the Genome Data Viewer (https://www.ncbi.nlm.nih.gov/genome/gdv/). The genomic structure of BF OT was predicted using genome sequences from lonStrDom2. Genomic regions surrounding OT were compared among human, rat, and BF using the Genome Data Viewer to analyze the synteny relationship.

For analysis of transcription factor binding sites, zPicture (https://zpicture.dcode.org/) and rVista (https://rvista.dcode.org/cgi-bin/rVA.cgi?rID=zpr09032019035446775) were used. zPicture alignments can be automatically submitted to rVista 2.0 to identify conserved transcription factor binding sites. RVista excludes up to 95% false-positive transcription factor binding site predictions, while maintaining a high search sensitivity.

To test for possible functional effects of the SNPs we identified, we used the Variant effect predictor tool, available in Ensembl (v. 100), for the respective sites in both the BF (LonStrDom1) and the chicken genomes (GRCg6a). Further, we used PhyloP (phyloP77way), available in the UCSC Browser, that scores the measure of evolutionary conservation at individual sites, by aligning 77 vertebrate species’ genomes. The scores are compared to the evolution that is expected under neutral drift, so that positive scores measure conservation, which is slower evolution than expected, while negative scores measure acceleration, which is faster evolution than expected^29^. Using this measure, we can infer whether some elements are functional, based on the rationale used widely in comparative genomics that functional genomic elements evolve more slowly than neutral sequences^30^.

We performed a multi-species alignment using CLUSTAL W (1.81), visualized via JalView (2.11.1.0) with the OT sequence of the following 29 avian species: BF, Zebra finch, Gouldian finch, Small tree finch, Medium ground-finch, Dark-eyed junco, White-throated sparrow, Blue tit, Rufous-capped babbler, Silver-eye, Flycatcher, Blue-crowned manakin, Eurasian sparrowhawk, Ruff, Spoon-billed sandpiper, Pink-footed goose, Swan goose, Golden pheasant, Ring-necked pheasant, Chicken, Indian peafowl, Turkey, Japanese quail, Helmeted guineafowl, Great spotted kiwi, Little spotted kiwi, Okarito brown kiwi, African ostrich and Chilean tinamou (for the IDs of the genomes used, Gene-IDs and locations used for the alignment: Supplementary Table 3). We used webPRANK (https://www.ebi.ac.uk/goldman-srv/webprank/) to confirm the ancestral state inference of the ‘GCT’ site in BF.

Further, a Maximum Likelihood phylogenetic amino acid tree was constructed for OT in all the avian species available in Ensembl (v.100), using TreeFAM and TreeBeST5 pipeline in the Ensembl ‘Gene tree’ tool package (https://www.ensembl.org/info/genome/compara/homology_method.html).

To test for intra-species variation of these sites in other avian species, we used the dbSNP (release 150) available for chicken (remapped to GRCg6a), dbSNP (release 139) available for turkey (Turkey_2.01), and dbSNP (release 148) available for zebra finch (bTaeGut1_v1.p).

### 2.5 ISH

We designed antisense probes to bind to both variant 1 and 2 of OT mRNA. Antisense probes for the OT mRNA were synthesized from the BF OT cDNA (512 bases, corresponding to variant 2 nt 1–512) with a ccc (3 bases, corresponding to variant 1 nt 510–512). Corresponding sense probes were also synthesized for control ISH (Suppl. Fig.S1). BF OT cDNA fragments were inserted into the pGEM-T easy vectors (Promega). The plasmids were digested with restriction enzymes (NcoI or SpeI) to release the fragment; probes were synthesized using SP6 or T7 RNA polymerase (Roche Diagnostics) with digoxigenin-labeling mix (Roche Diagnostics).

Frozen brains were cut in 20-μm coronal sections on a cryostat (Leica Microsystems, Wetzlar, Germany). Every thirteenth section through the hypothalamus from each animal was mounted on 3-aminopropyltriethoxysilane-coated slides and stored at −80°C until use. ISH was performed as described previously^31^, except that sections were postfixed in 4% paraformaldehyde for 10 min, proteinase K treatment was omitted and color development of alkaline phosphatase activity was carried out for 1 h with nitroblue tetrazolium chloride (Roche Diagnostics) and 5-bromo-4-chloro-3-indolyl phosphate (Roche Diagnostics) in detection buffer. Images of sections were captured with a LeicaMC170 HD camera attached to a Leica DM500 microscope (Leica Microsystems).

### 2.6 Analysis of OT mRNA expressing cells

The “analyze particle” module of ImageJ (based on size = 50-infinity pixel and threshold =0-150) was used to count the numbers of OT mRNA-expressing cells in the following regions of interest: the lateral hypothalamus (LHy) on the left side of the brain; the external subgroups of the supraoptic nucleus (SOe) on the left side of the brain; and PVN at the level of the anterior commissure on the both sides. This limitation was due to the absence of the lateral two-thirds of the right brain hemisphere of a WRM. The sum of cell numbers in each area was calculated.

Statistical analyses for OT mRNA expressing cells

Results are shown as means ± standard errors of the mean and coefficients of variation. Statistical analyses were conducted with Prism 4.0 (GraphPad Software, USA), using unpaired Student’s t-tests. P values <0.05 were considered statistically significant.

### 2.7 Real-time qPCR

We isolated RNA from each different brain region. Total RNA was extracted using QIAzol Lysis Reagent and column purified (RNeasy Lipid Tissue Mini Kit: Qiagen). We performed reverse transcription using Superscript First-Strand Synthesis (Invitrogen) with oligo (dT) primer. Primers for OT were designed using sequences downloaded from GenBank or Ensembl (Supplementary Table 4), and for the OTR were designed based on previous studies^32,33^. The control gene was peptidylprolyl isomerase A (PPIA), which have been evaluated in two species of songbird: zebra finch and white-throated sparrow as highly stable in the brain^34^. qPCR was performed using Roche LightCycler 96 System with TB Green Premix Ex Taq II (Takara), in triplicate for each sample on 96-well plate. We calculated crossing point values (CT) using the Abs Quant/ 2nd Derivative Max method using LightCycler 96 SW 1.1 software. CT was used to calculate ΔΔCT ([CT target gene-CTcontrol gene] - [CT target gene-CTcontrol gene] calibrator). We picked average CT of BF samples as calibrator. Relative expression levels between the species were calculated using the 2-ΔΔCT method^35^.

### 2.8 Production of antiserum to the avian OT

The antiserum was raised in a rabbit immunized 5 times with synthetic polypeptide of the avian OT (mesotocin) sequence (CYIQNCPIG-amide) as antigen every 1 weeks. The specificity of the serum was tested by a dot immunoblot assay, with the OT and its homolog and paralog (vasotocin) in other species (OT, isotocin, vasotocin, vasopressin), where we confirmed it binds solely to avian and fish OT (isotocin). Aliquots of the homologs (oxytocin, mesotocin, isotocin) and paralogs (vasotocin, and vasopressin) peptides were spotted onto a 0.2-μm polyvinylidene fluoride membrane (Immun-Blot PVDF Membrane for Protein Blotting, BIO-RAD). The membrane was air dried at room temperature and was washed for 10 min in 0.05 M Tris buffer (pH 7.6) with 0.1% Tween 20 and 0.15 M NaCl (TBS-T) and incubated for 60 min in blocking solution containing 5% skim milk in TBS-T. After blockage, the membrane was exposed for 120 min to mesotocin antiserum (1:1,000 dilution in blocking buffer). After the primary immunoreaction, the membrane was further incubated with anti-rabbit biotinylated IgG secondary antibody (Agilent Technologies, Inc., Santa Clara, CA, USA; 1:200 dilution in blocking buffer) for 120 min and Avidin-Biotin Complex Reagent (Vector Laboratories, Inc., Burlingame, CA) for 120 min, and subsequently with ImmPACT DAB peroxidase substrate solution (Vector Laboratories) for 5 min at room temperature.

### 2.9 OT EIA

The frozen brain tissue blocks of WRM (n = 7) were dissected as described above immediately before peptide extraction. The frozen brain tissue blocks of WRM and BF were boiled for 8 min and homogenized in 5% acetic acid using a TissueLyser LT (Qiagen) for 6 min at 50 Hz. The homogenate was centrifuged at 14,000 rpm for 30 min at 4°C. The supernatant was collected and forced through a disposable C-18 cartridge (Sep-Pak Vac 1cc; Waters, Milford, MA, USA). The retained material was then eluted with 60% methanol. The pooled eluate was concentrated in a vacuum evaporator, passed through disposable Ultrafree-MC centrifugal filter units (Millipore, eBillerica, MA, USA) and dried. The dried material was reconstituted in 220 μl Dilution buffer, and 100 μl of the sample was used for EIA at duplicate.

The samples were subjected to competitive EIA by using the antiserum described above. In brief, different concentrations of OT (0.01-100 pmol) and adjusted tissue and plasma extracts were added with the antiserum against OT (1:1,000 dilution) to each antigen-coated well of a 96-well microplate (F96 maxisorp nunc-immuno plate;Thermo Fisher Scientific, Roskilde, Denmark) and incubated overnight at 4°C. After the reaction with alkaline phosphatase-labelled goat anti-rabbit IgG (Sigma; 1:1,000 dilution in dilution buffer), immunoreactive products were obtained in a substrate solution of p-nitrophenylphosphate (SIGMA FAST™ p-nitrophenyl phosphate tablet set) for 90 min, then, 20 μL 5 N NaOH solution was added to and the absorbance was read at 405 nm with a reference filter of 620 nm by iMark Microplate Reader S/N 20255 (Bio-Rad, USA).

### 2.10 Statistical analyses for qPCR and EIA

For analysis of the EIA results, statistical analyses were conducted with Prism 4.0 (GraphPad Software, USA), using F-tests, and unpaired Student’s t-tests with Welch’s correction. For analysis of the qPCR results, we performed repeated measures ANOVA (species and brain region) followed by Holm-Bonferroni correction for multiple comparison using statistical software R (http://www.r-project.org/; R Core Team, 2013; R Foundation for Statistical Computing) to investigate the difference of relative gene expression levels between BF and WRM. P values <0.05 were considered statistically significant.

## 3 Results

### 3.1 Identification and comparison of *OT* cDNA in Bengalese finch and white-rumped munia

We performed PCR amplification with primers, based on the available predicted zebra finch *OT* cDNA in the Ensembl genome browser (version 98.1), and isolated *OT* transcripts from brain tissues of five BFs s and four WRMs (Suppl. Data 3). We found that two of the five BF shared an identical *OT* transcript, which we named ‘variant 1’, while the transcript we isolated from the other three BF (‘variant 2’) showed specific changes in the 5’- and 3’-UTRs with respect to variant 1. Specifically, variant 1 contained a 3-nucleotide (GCT) deletion in the 5’-UTR, and a single nucleotide polymorphism (T > C transition) in the 3’-UTR (Fig. 1A). To confirm that our sequence is the BF *OT*-ortholog, we BLAST searched the full-length cDNA sequence against a reference BF genome assembly (lonStrDom2; accession GCF_005870125.1)^36^. Only one locus (BLAST hit) in the genome showed high similarity (E-value < 5e-56) that was unnamed (ID: LOC110473283), located on chromosome 4 (68,957,139-68,958,557). This sequence was identical to variant 2. Combining genotypes, we confer that four out of the six *OT* sequences are of the variant 2-type, and given that the reference genome is from an University of California laboratory population separate from the Japan laboratory population, this indicates that variant 2 might be the dominant variant for the strain. All four *OT* transcripts we isolated from WRM did not show any variability, and were identical to the variant 1 of the BF, except for a single nucleotide change in position 416, where WRM had A and all six BF-sequences had G in the 3’-UTR (Fig. 1A). Otherwise, they were identical in the 369-nucleotide protein coding sequences (Fig. 1A), the open reading frame (ORF)-region that encodes the 19-amino-acid signal peptide (Fig. 1A, underlined), the 9-amino-acid nonapeptide *OT* hormone (black), the tripeptide processing signal (GKR), and the 92-amino-acid neurophysin I sequence (gray).

**Fig. 1.**
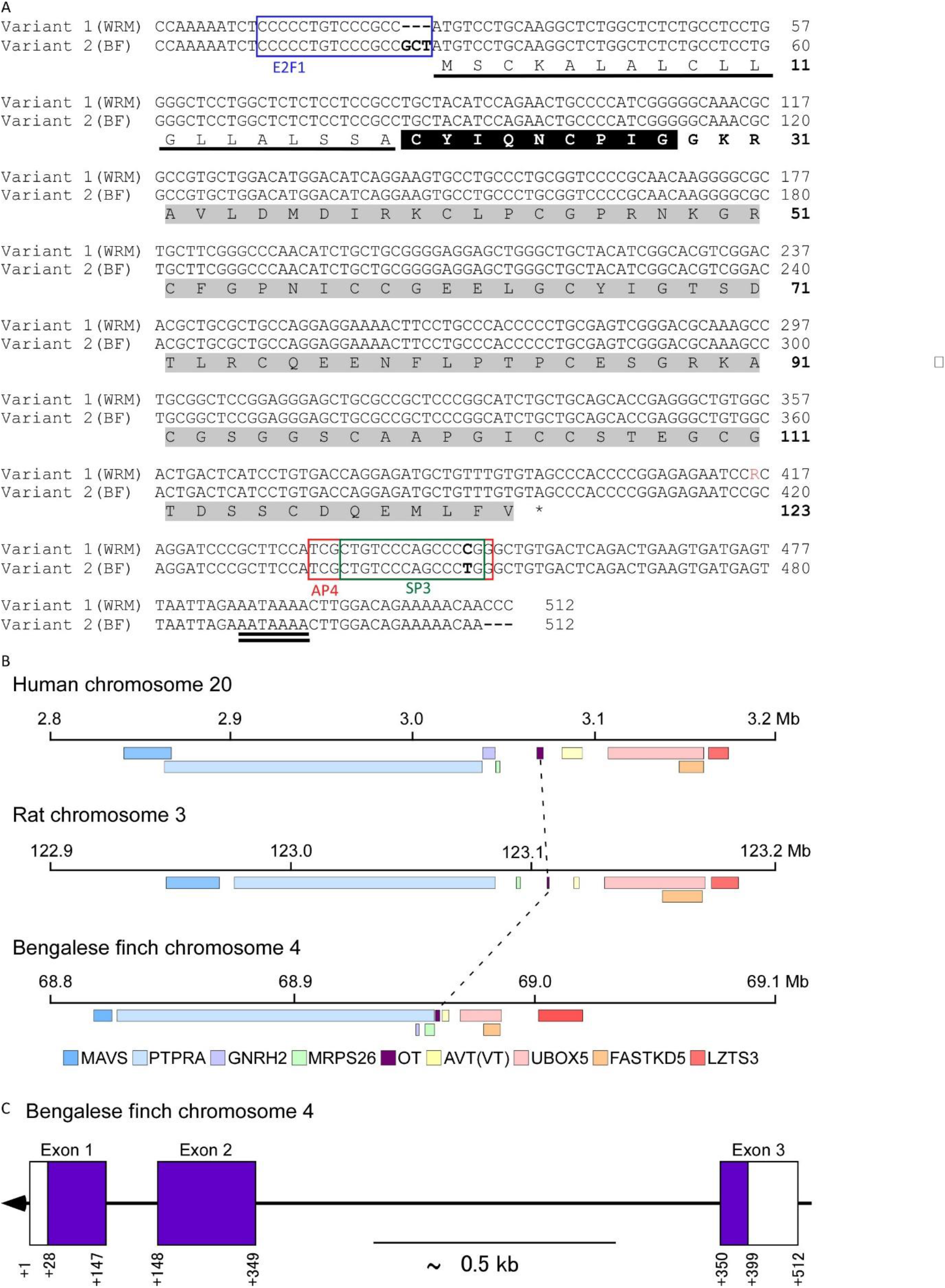
Nucleotide and amino acid sequences of white-rumped munia and Bengalese finch oxytocin (*OT*) precursor cDNA. (A) *OT* sequence is on a black background. Neurophysin I sequence is on a gray background. Tripeptide processing signal (GKR) is indicated in bold. Predicted signal peptide is underlined. Poly (A) adenylation signal AATAAA is indicated by double-underline. Sequences of putative transcription factor binding sites are boxed. Differences in nucleotide sequences are indicated with bold letters. AP4, activating enhancer binding protein 4; E2F1, E2F transcription factor 1; SP3, specificity protein 3. (B) Schematic representation of Bengalese finch *OT* structure. Exons are boxed and numbered, and introns appear as straight black lines. Shaded and open boxes denote coding and noncoding sequences, respectively. (C) Conserved genome synteny among human, rat, and Bengalese finch *OT* loci. Human, rat, and Bengalese finch genomic organizations were obtained by using Genome Data Viewer. Genes are indicated by shaded boxes. *OT* genes are linked by dotted lines. In rodents, gonadotropin-releasing hormone 2 gene (GnRH2) is inactivated or deleted from the genomes^65^. WRM, white-rumped munia. BF, Bengalese finch.

### 3.2 Synteny and phylogenetic analyses confirm our identified gene in BF as the *OT*-ortholog

To further confirm our identified gene in the BF was the true *OT*-ortholog, we ran synteny analysis on the surrounding territory of the identified gene in BF (LOC110473283). The analysis revealed that this gene is located in a genomic region that is highly conserved in tetrapod vertebrates, exactly where the *OT* is found in those species (Fig. 1B). Conserved syntenic genes in the surrounding territory include mitochondrial antiviral signaling protein (*MAVS*), protein tyrosine phosphatase receptor type A (*PTPRA*), mitochondrial ribosomal Protein S26 (*MRP26*), arginine vasopressin/vasotocin (*AVP/VT*), fAST kinase domains 5 (*FASTKD5*), U-box domain containing 5 (*UBOX5*), and leucine zipper tumor suppressor family member 3 (*LZTS3*) (Fig. 1B). Interestingly, in the BF genome (lonStrDom2), *AVP/VT* was also unnamed (Gene ID: LOC110473284). Based on our synteny analysis, and on further synteny and phylogenetic results presented in Theofanopoulou et al. (accepted)^37^, we propose that LOC110473283 should be named oxytocin (*OT*) and LOC110473284 should be named vasotocin (*VT*) in the BF. Our phylogenetic tree using the BF putative *OT* in LonStrDom1 as a query sequence (ENSLSDG00000001590; unnamed) grouped this sequence with the *OT*-ortholog in all vertebrates (Suppl. Fig. S2 See Gene Tree ID: ENSGT00390000004511 for a capture of the tree only in avian species). These phylogenetic results corroborate with our synteny results in that the gene we identified in the BF is the true *OT*-ortholog.

Using the *OT* sequence from lonStrDom2 (chr4: 68,957,139-68,958,557) and our identified BF *OT* cDNA sequence, we further predicted the *OT* genomic structure (Fig. 1C). The BF *OT* consists of three exons, spanning 1,419 base pairs of genomic sequence. Exon 1 contains the 5’-UTR, signal peptide, *OT*, nonapeptide (CYIQNCPIG), and a portion of the neurophysin peptide sequence. Exon 2 and the first 48 base pairs of exon 3 contain the remaining sequence of the neurophysin peptide; the remainder of exon 3 contains the 3’-UTR (Fig. 1C).

### 3.3 Transcription factor-binding and functional analyses on the identified changes

Our analysis using zPicture and rVista softwares showed that the 5’-UTR GCT deletion is located at the end of a putative binding site of the *E2F1* transcription factor (Fig. 1A, blue box). The 3’-UTR C > T transition is located in a putative binding site of the AP4 and SP3 transcription factors, upstream of the polyadenylation signal (aaataaa). These transcription factor binding sites (TFBS) were present only in the BF and WRM, but were not conserved in other species we tested, like chicken, zebra finch and human.

Variant effect predictor test that predicts functionality of specific single nucleotide polymorphisms (SNPs), applied to the respective sites in both the BF (LonStrDom1) and the chicken genomes (GRCg6a) did not give any known or novel functional effect of these sites. A PhyloP test that scores evolutionary conservation at individual sites in a 77 vertebrate species’ genome alignment (phyloP77way), showed that the A > G transition site had a mean positive score (1.02034; Suppl. Fig.S3A), while the C > T transition site had a mean negative score (−0.0770591; Suppl. Fig. S3B). This suggests that the A > G site is a more evolutionary conserved, hence a more functional site, based on the view that conserved sites across species tend to be functional.

### 3.4 Comparison of *OT* mRNA sequence in avian species

To determine if the changes we observed in the variant 2 of *OT* were specific to BF or shared with other bird species, we compared it with *OT* sequences from 29 avian species (Fig. 2, Suppl. Fig. S2, Supplementary Table 3). Surprisingly, a 5’-UTR deletion was found only in WRM. However, different species had different sequences: GCT was only present in the Gouldian finch, the Silver-eye and the Blue tit, apart from the BF, while most species (22) had a GCC. Specifically, G (first nucleotide of the deleted trinucleotide sequence; GCT in the WRM) is conserved across 93% of the species, while 7% of the species in the alignment show A; C (second nucleotide of the deletion) is present in all avian species studied except WRM (99%); and T (third nucleotide of the deletion) is present in only 21% of the birds, with 76% having C on this site (A: 3%). As these percentages suggest and as we confirmed via Webrank, GCC constitutes the ancestral state of this site, which was lost in WRM and some BF.

**Fig. 2.**
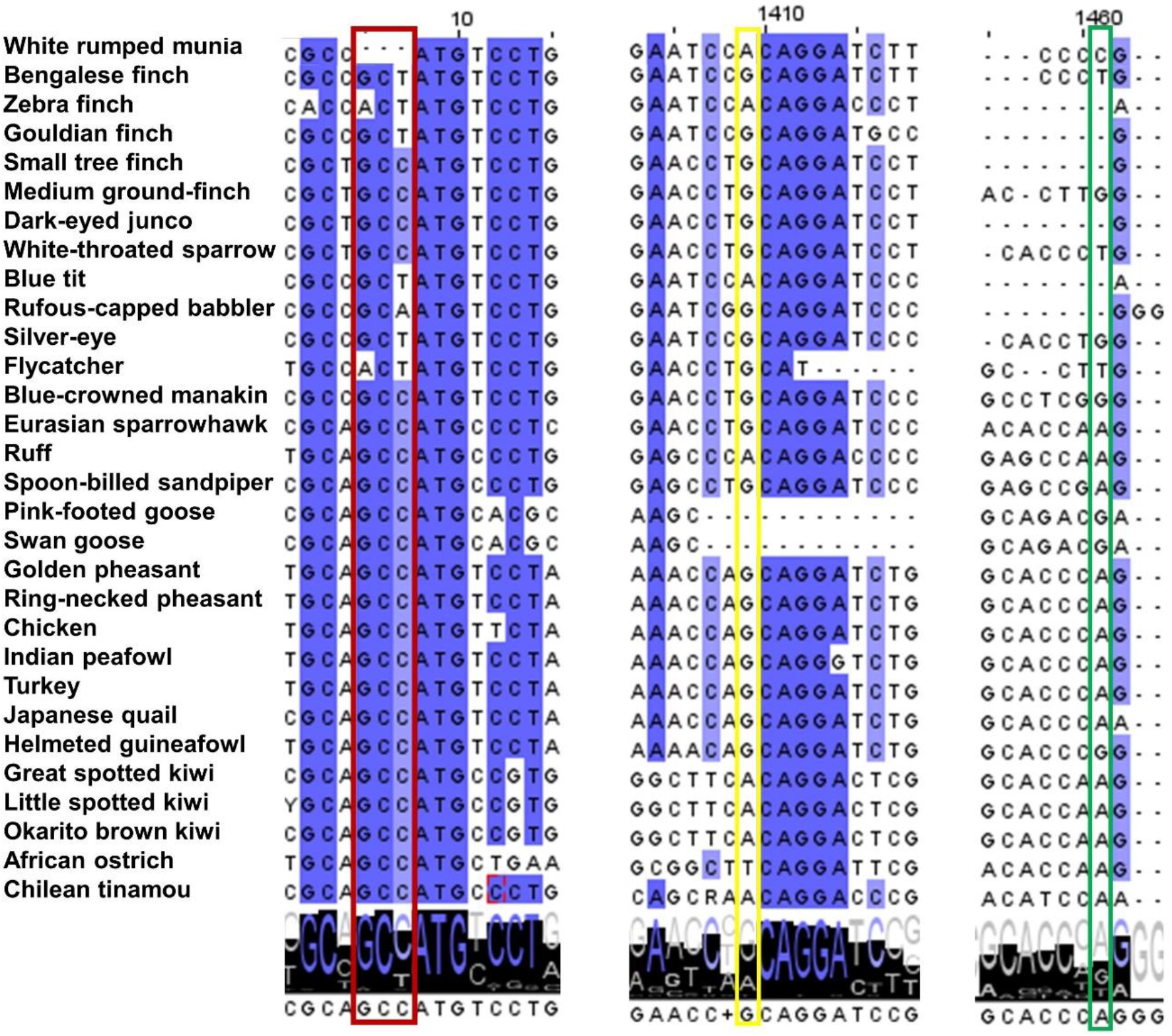
Multi-species alignment of the oxytocin (*OT*) in 30 avian species, with specific focus on the three sites that differ in BF and WRM. 1^st^ column: GCT deletion site in WRM, 2^nd^ column: A > G transition site, 3^rd^ column: C > T transition site. Alignment was made using ClustalW, visualized through JalView.

In the 3’-UTR, in the A > G transition site (2^nd^ column of Fig. 2), G is present in most of the avian species of our alignment (66%) (A: 24%; T: 3%), while the C > T transition site (3^rd^ column of Fig. 2) is more variant across species (A: 48%; G: 21%; T: 10%); in addition to the BF, only the White-throated sparrow and the Flycatcher had T in this site. These results corroborate and extend our PhyloP data from the alignment of 77 vertebrates, and also show that in avian species the A > G site is more conserved than the C > T site. Although this suggests a higher predicted functionality for the A > G site, the T allele (in the C > T transition site) is rare in birds (10%), something that suggests that this variant might be functional in the avian lineage.

Among the avian species we used in our alignment, some of them come from domesticated species (chicken, zebra finch, turkey, Japanese quail and swan goose), but we were not able to find any pattern of convergent evolution among them. Only the G allele (A > G site) was shared by the BF, the chicken, the quail, the swan goose and turkey, but we do not consider this evidence of convergence since G is present in most avian species (66%) of our alignment. We further tested for possible intra-species variation of these sites in other avian species, using the dbSNP (release 150) available for chicken (GRCg6a), dbSNP (release 139) available for turkey (Turkey_2.01), and dbSNP (release 148) available for zebra finch (bTaeGut1_v1.p), but we did not find any variation in the sites studied within any of these species.

### 3.5 Changes in *OT* mRNA expression in the diencephalon of BF relative to WRM

Differential *OT* gene expression in several diencephalic nuclei has been associated with social behavioral differences in several species^38^. Since *OT* expression is highly restricted to the hypothalamus within the brain^39,40^, we performed ISH for *OT* mRNA in BF and WRM brain sections containing several hypothalamic nuclei. BF and WRM had similar distributions of *OT* mRNA-expressing cells in the hypothalamus (Fig. 3, Suppl. Fig.S1 for ISH with a sense and antisense OT probe), and we did not detect any significant differences in the numbers of cells expressing *OT* mRNA (Fig. 4A-C, Supplementary Table 5) in any of the nuclei tested between the two strains by ISH. These nuclei included the SOe, LHy, and PVN (Fig. 3A-F). We found that the PVN contained a higher average number of *OT* cells in both strains (48.0 in BF; 46.7 in the WRM), compared to the SOe (7.4 in BF; 11.3 in the WRM) and the LHy (7.0 in BF; 7.6 in the WRM) (Supplementary Table 5).

**Fig. 3.**
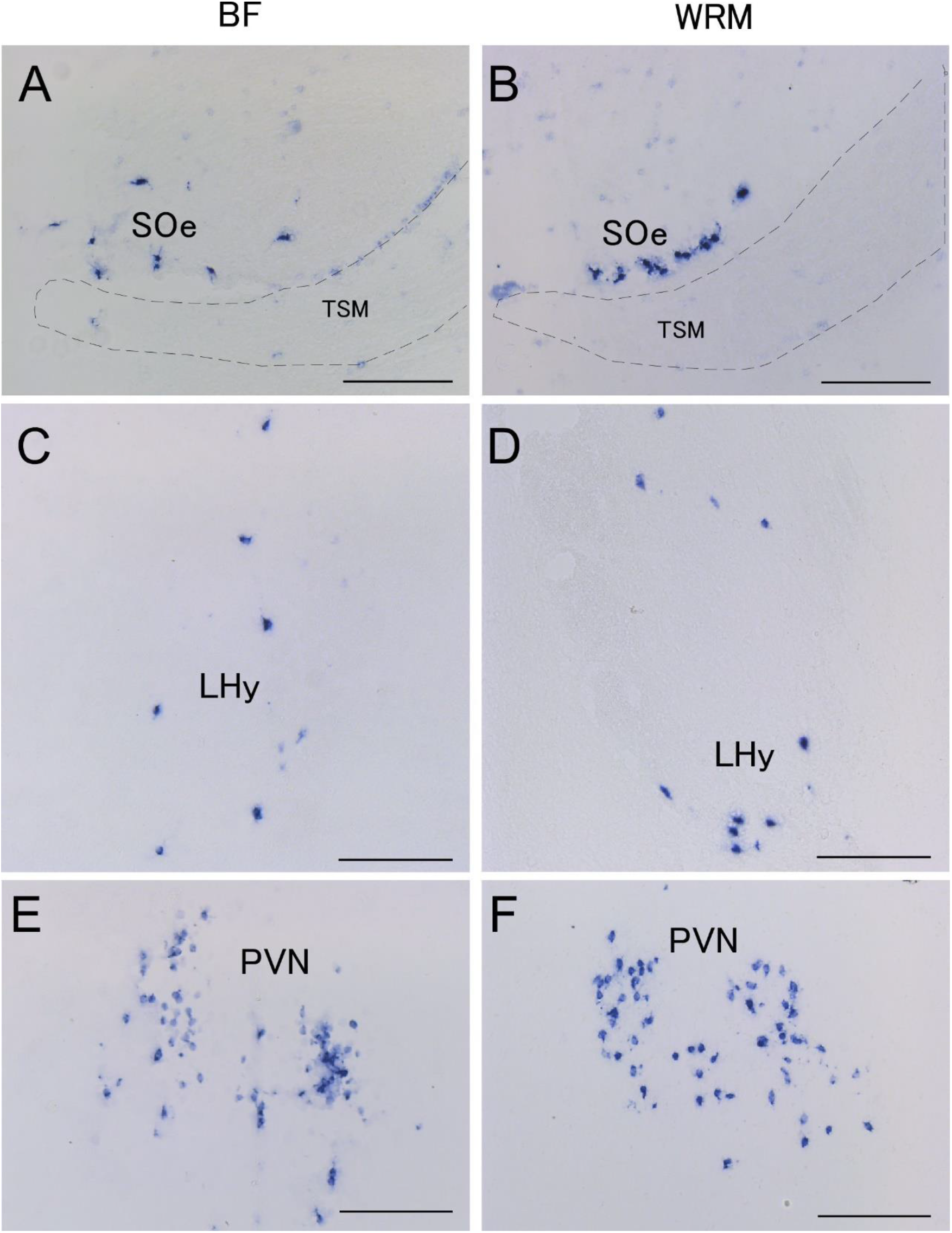
Photomicrographs of oxytocin (*OT*) mRNA-expressing cells in brains of Bengalese finches and white-rumped munias. (A, B) SOe, external subgroups of supraoptic nucleus. (C, D) LHy, lateral hypothalamic areas. (E, F) PVN, paraventricular nucleus. TSM, septopalliomesencephalic tract. Scale bars = 200 μm.

**Fig. 4.**
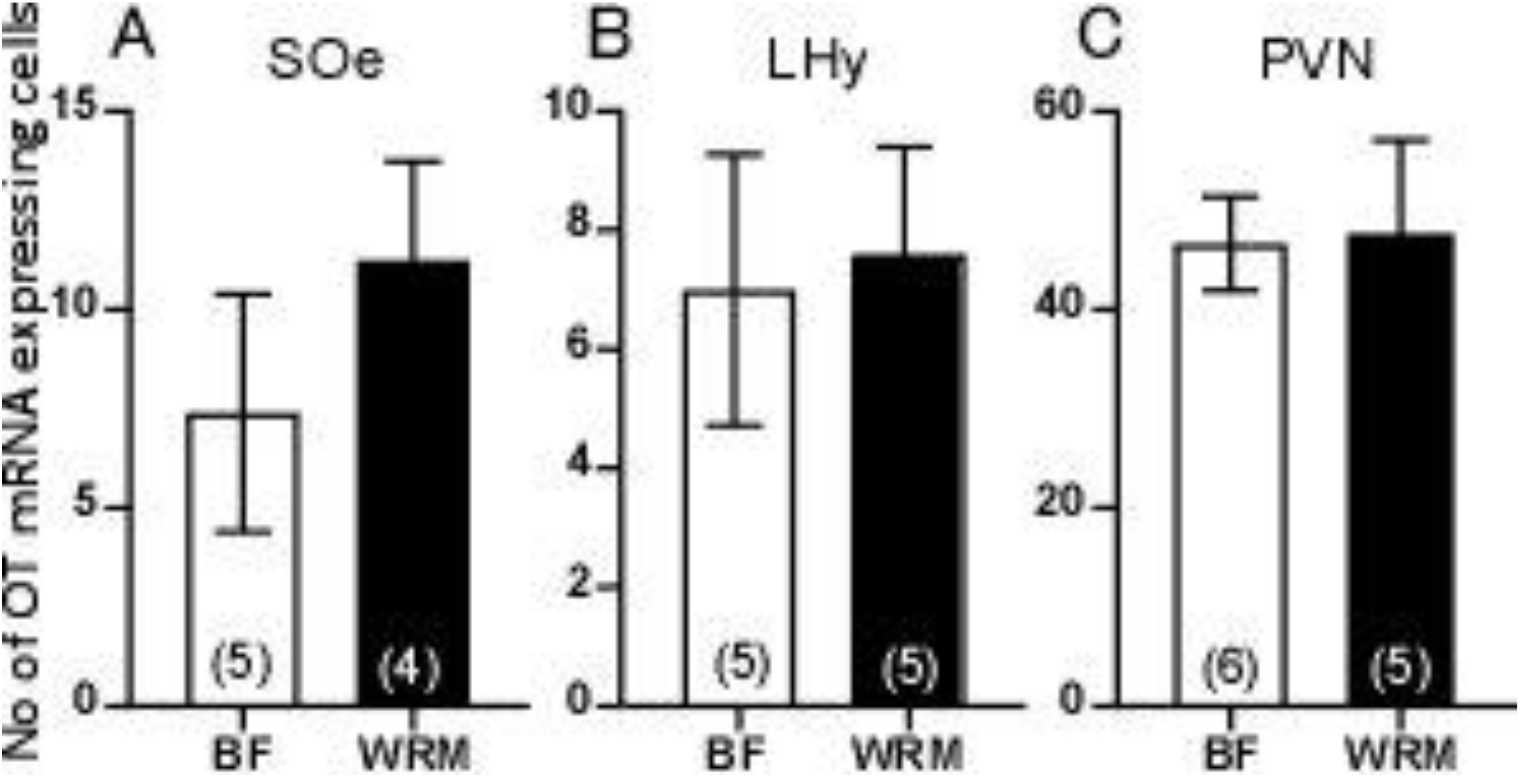
Comparisons between the number of cells expressing OT in hypothalamic nuclei in the Bengalese finches and white-rumped munias. (A) External subgroups of the supraoptic nucleus (SOe). (B) Lateral hypothalamus (LHy). (C) Paraventricular nucleus (PVN). Numbers in bars denote numbers of birds analyzed. Data are means ± standard errors.

To compare mRNA expression levels, we performed qPCR to measure relative *OT* mRNA expression in the BF and WRM cerebrum and diencephalon (which contains the hypothalamus). We found that *OT* mRNA expression of BF in the diencephalon was significantly lower than that of WRM (repeated measures ANOVA followed by Holm-Bonferroni correction, F(1,10) = 12.2243, p = 0.0058, Fig. 5, Supplementary Table 6), while we found no significant differences in the cerebrum (repeated measures ANOVA followed by Holm-Bonferroni correction, F(1,10) = 3.9622, p = 0.0746, Fig. 5).

**Fig. 5:**
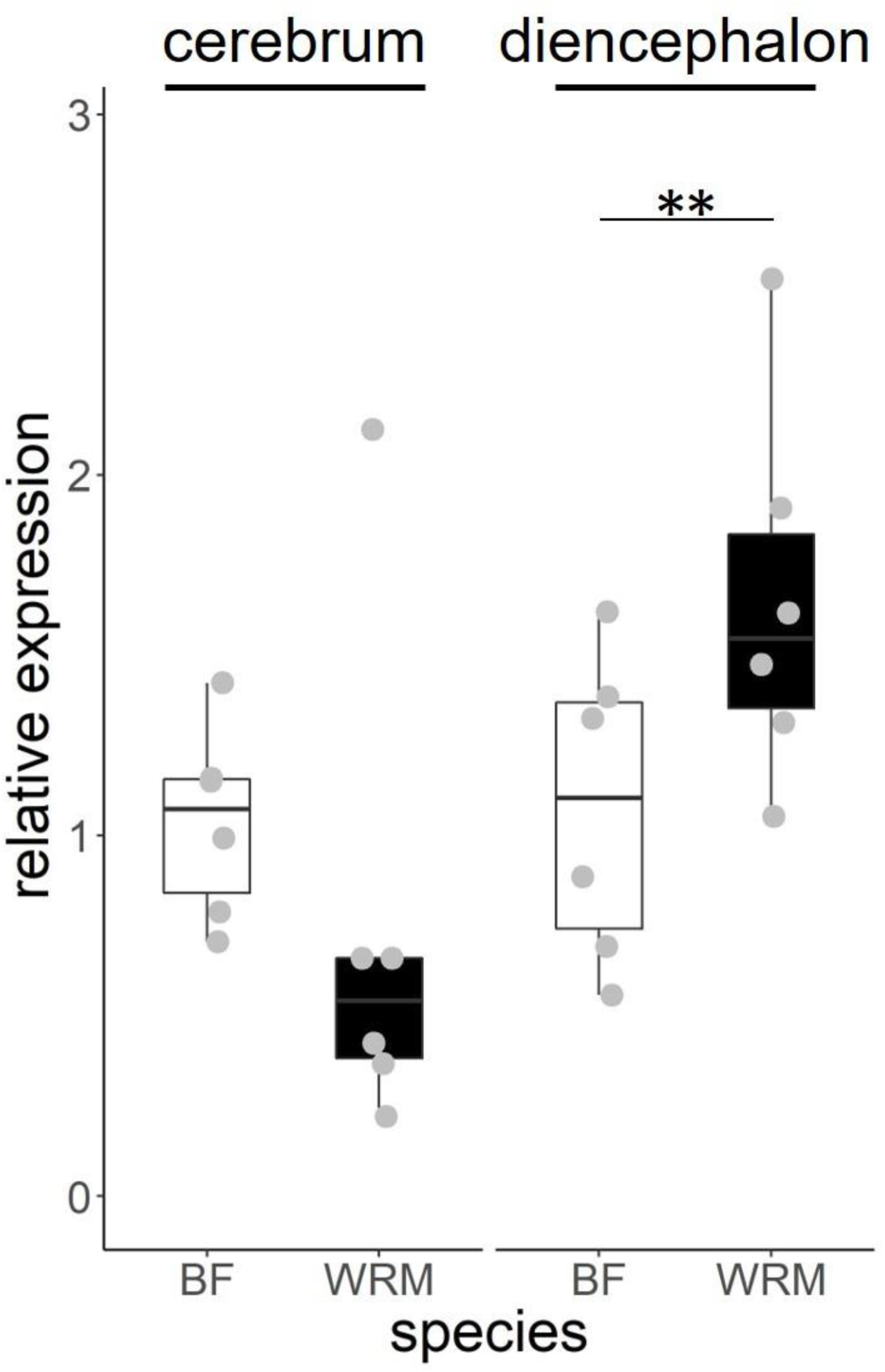
Cerebral and diencephalic oxytocin expression in Bengalese finches and white-rumped munias. Each dot indicates average of individual value (BF: n = 6, WRM: n = 6). **p < 0.01.

To investigate whether the difference of *OT* synthesis in the diencephalon of BF and WRM affects *OTR* expression, we conducted qPCR quantifying the levels of *OTR* mRNA in the cerebrum, diencephalon, midbrain and cerebellum, but we did not detect any significant differences (repeated measures ANOVA, species: F(1,10) = 0.1693, p = 0.6894, species × brain region: F(3, 30) = 0.6394, p = 0.5956, Supplementary Table 7, Fig. 6).

**Fig 6:**
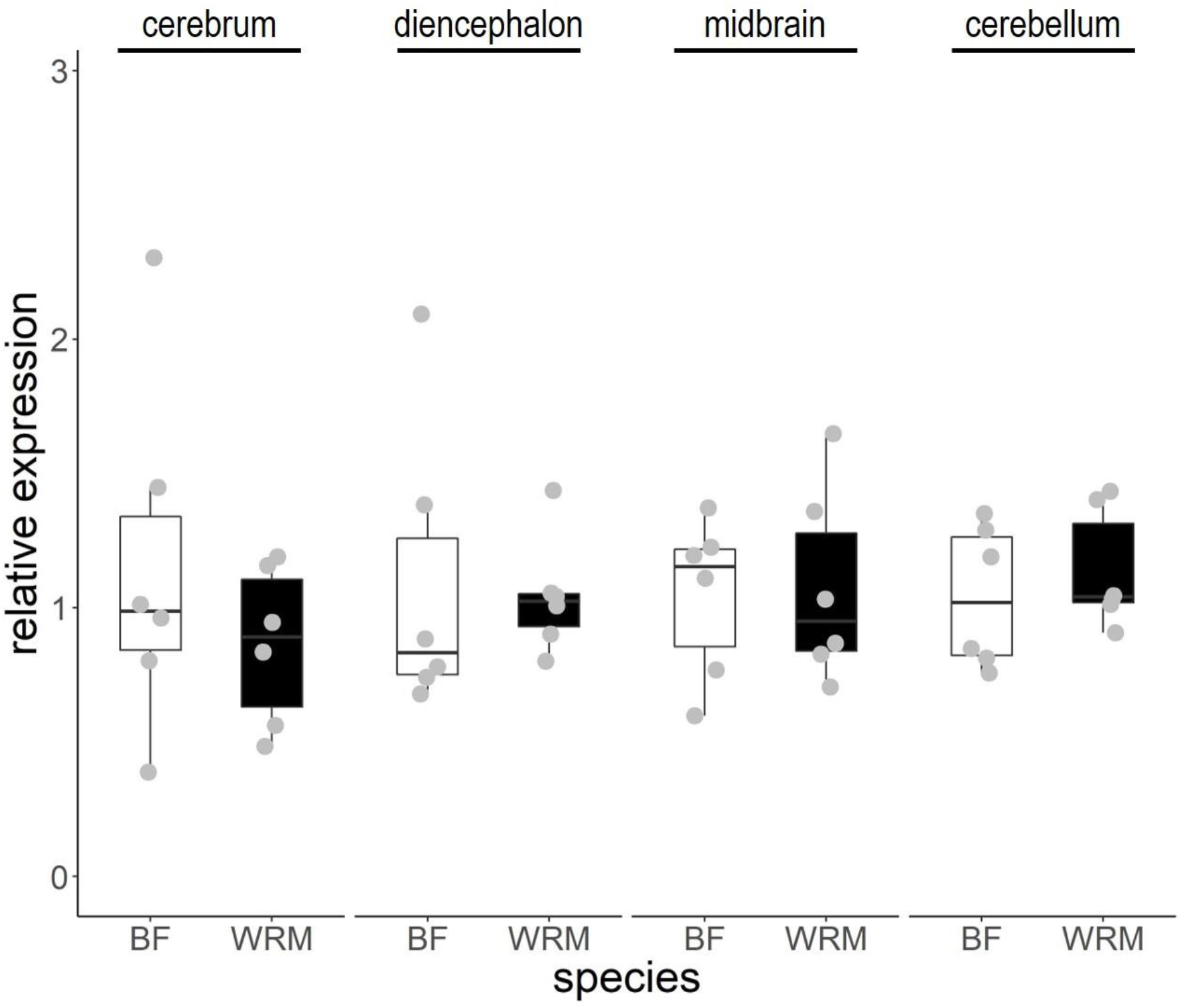
Oxytocin receptor (*OTR*) expression in different brain regions in Bengalese finches and white-rumped munias. Each dot indicates the average of individual value (BF: n = 6, WRM: n = 6). *OTR* exhibited no significant differences in relative expression levels between WRM and BF in these brain areas.

We generated a specific antiserum against the avian *OT* to quantify *OT* brain peptide levels in both strains. Our *OT* antiserum bound to the avian and fish *OT* peptide (Fig. 7A), but did not cross-react with either the mammalian *OT* or with *VT* of any species. We lastly compared brain *OT* peptide levels using EIA (Fig. 7B; Supplementary Table 8) in both strains. Although we did not detect significant differences between strains (t = 1.630, df = 6.041, p = 0.1539), we found a significantly higher intra-strain variability in the WRM compared to the BF (F = 293.1, Dfn = 6, Dfd = 6,*p* < 0.0001). However, consistent with the qPCR results, OT peptide content levels in the brains of BF were more clustered at low levels than those of WRM.

**Fig. 7:**
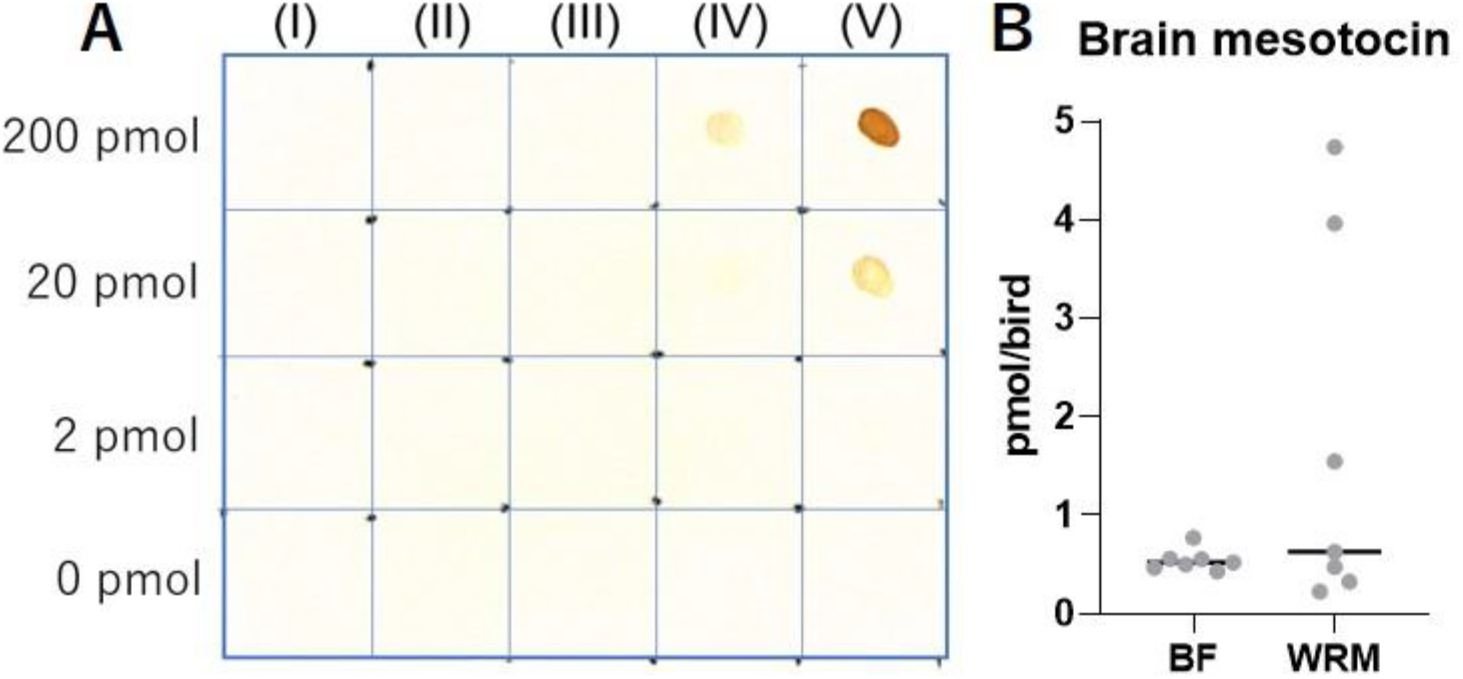
Dot immunoassay demonstrating antiserum specificity to avian and fish oxytocin (*OT*) and brain oxytocin content of Bengalese finches and white-rumped munias. (A) From 0 to 200 pmol of mammalian vasotocin (I), avian vasotocin (II), mammalian oxytocin (III), avian oxytocin, (IV) or fish oxytocin (V), peptides were spotted onto a polyvinylidene difluoride membrane. The membrane was incubated in avian oxytocin antiserum (1:1,000 dilution) and the immunoreaction was visualized using biotinylated secondary antibody and avidin-biotin complex. (B) WRMs exhibit larger individual variation in the brain OT content compared to BFs. Each dot indicates individual value, and the vertical line indicates median.

## 4 Discussion

In this article we sought to identify whether differences in the sequence and brain expression of *OT* in the wild WRM and the domesticated BF would reveal insights in the neuroendocrinological changes that underlie the domestication process of the latter. We found two different genotypes for *OT* in BF (variant 1 and variant 2) differing in the 5’-UTR (GCT) and 3’-UTR (C > T) sequences. The *OT* we isolated from the WRM was identical to the BF variant 1, except for an additional change in the 416^th^ position (A > G). Thus, the WRM *OT* differs from both BF-variants in the A > G change, and to BF-variant 2 in particular, it also differs in the 5’-UTR GCT deletion and the 3’-UTR C > T change.

Although our BF and WRM sample size is too small to make predictions of selection pressures, the inter-strain variation findings help define what is special. Only the WRM and some BF had the 3 nucleotide 5’-UTR deletion, indicating that this occurred among the two strains before BF was domesticated, considering that in other species we searched for in their SNP databases (chicken, turkey, zebra finch), there were no variation patterns on the *OT* sites that showed variation between WRM and BF. Although this variation can be suspected to come from laboratory population bottlenecks, in fact, 3 of the 4 ‘variant 2’ individuals come from the colony at RIKEN, Japan while the remaining 1 individual comes from a different UCSF colony in California. Thus, this variation does not seem to be due to laboratory population bottlenecks.

The absence of nucleotide change convergence in BF and other domesticated avian species should not come by surprise as domestication processes have been shown to not obligatorily work through selection on the same site, but also through selection on the same gene or biological process^41^. Nonetheless, we do not exclude the possibility that convergent evolution on different nucleotides in the same gene can occur.

Our findings that the 5’-UTR GCT variant is located at the end of a putative binding site of the *E2F1* transcription factor (Fig. 1A, blue box), and the 3’-UTR C > T in a putative binding site of the AP4 and SP3 transcription factors indicate that they may affect mRNA regulation. Such changes could affect binding sites, thereby modifying the transcription rate, subcellular localization, mRNA stability, and/or synthesized protein quantity^42–45^. Although we did not find these transcription factor binding sites (TFBS) being conserved in the *OT* of the others species we studied (chicken, zebra finch, human), which would traditionally be viewed as a weaker predictive evidence, an array of recent studies have shown that most TFBS are species-specific, and aligned binding events present in many species are rare^46,47^. Thus, these TFBS could be newly derived elements, specific to BF and WRM, if not in other closely related species.

Our ISH findings largely agree with previously identified *OT* distributions in other avian species (including in chicken^39^, zebra finch^40,48^, Japanese quail^48,49^, domestic mallard^48,49^ and starling^48,49^), in that most of the OT-containing cells we identified were in the PVN, compared to the other nuclei we tested (SOe, LHy) and the rest of the brain. Vicario et al. (2017)^40^ found age-dependent downregulation of *OT* in the PVN of zebra finches, indicating that the expression levels can change over time within a species. Lastly, it has been shown in chicken^39^ that some *OT* cells in the PVN could also express *VT*, but in the domestic duck and the Japanese quail^49^, *OT* neurons are separate from *VT* neurons in the PVN. Future investigations in the munia finch and BF will need to be done to determine if there is or isn’t any overlap in the two sister genes.

The significantly lower amount of *OT* mRNA in the diencephalon of the BF compared to WRM is intriguing in that it implies there is less *OT* synthesis that could be associated with domestication. Our finding of less *OT* synthesis in the domesticated BF is in the opposite direction from those reported for brain *OT* synthesis in rats and mice (laboratory domestication)^15^, where they found more *OT* production for the domesticated strain. This suggests that both higher and lower OT can be associated with domestication in different species or lineages. This could be due to these species (rats and mice vs. BFs) following a different path to domestication. Namely, both rats and mice underwent a process of human acclimation as companion animals during the 17th and 18th century, before they were used as laboratory animals^50,51^, while the BF did not undergo such an active process of human acclimation through the process of their domestication.

Concerning the mechanisms through which *OT* could be acting differently in the brain circuits of BF and WRM, one hypothesis is that *OT* from the hypothalamus would impinge differentially on neurons containing *OTR* in brain regions involved in social cognition and aggression regulation (e.g. striatum and lateral septum^25^). Such changes could account for the decreased aggression and fear seen in the BF, compared to the WRM^1,2,6^. Another possible brain mechanism could be more indirect, through the action of *OT* to the HPA axis. *OT* has been shown to inhibit the general reactivity of the HPA axis^52^, and to attenuate ACTH secretion and CORT levels^14^. More mechanistically, OT has been found to inhibit synaptic glutamate transmission onto CRH neurons, suppressing CRH neuron excitability and stress axis activity^53^. This mechanism could explain the lower CORT levels seen in BF^2^, along with the behavioral outcomes that this reduction implies.

A third possible mechanism, which also takes into account the reduced *OT* brain expression in BF, comes from studies in other avian species: In male violet-eared waxbills (*Uraeginthus granatina*), robust transcriptional activation of *OT* neurons in the PVN is observed following pursuits by a human hand^54^. *OT* neurons in the PVN also promote a passive stress-coping behavior in male zebra finches^55^. If the same mechanism applies in WRM, higher levels of *OT* in the PVN can explain why WRM shows a longer tonic immobility response following rapid inversion and restraint, compared to BF^1^. The tonic immobility response is an innate fear and defensive response associated with intensely dangerous situations^56^. This response is characterized by a temporary state of profound inactivity and relative lack of responsiveness to external stimuli^57^. The high level of *OT* may be needed to induce a longer tonic immobility reaction that reduces the threat of a potential predator and increases the chances of survival of WRM during a predatory attack in the wild. BF may no longer require high *OT* levels in the PVN to cope with predation because they have adapted to low predation pressures by domestication. Future experiments of *OT* manipulations in the PVN would help test this hypothesis.

In contrast to *OT* expression levels, we did not detect any significant differences in the expression of *OTR* mRNA in cerebrum, diencephalon, midbrain and cerebellum. This suggests that downregulation of *OT* in BF does not affect its receptor expression. OTR distributions in the telencephalon including the septum correlates with species-typical group size (gregarious or territorial) in estrildid finch species^58^. WRMs in Taiwan move and forage in flocks of around four to twenty birds^59^. Group size of BFs is difficult to define, but in our laboratories at the University of Tokyo and RIKEN, birds are kept in cages of similar flock sizes, and they are socially gregarious. It is possible then that the similar OTR distributions in both strains correlate with similar group sizes, as in other finch species, although we cannot rule out the possibility that strain differences in *OTR* expression could be detected using other experimental designs (e.g., larger sample sizes, conditions of breeding and sample collection, or quantifying mRNA level in specific brain regions by ISH or captured by laser microdissection). Other brain regions could include the song learning nuclei. *OTR* expression has been identified in vocal learning nuclei, like the HVC and RA^32,33,^, and based on different manipulations of the *OT* system, there is evidence that blocking the *OT* or the *OTR* can impact song learning (using the Manning Compound that blocks both *OTR-VTRs*^60^), or singing^9,26,61^. Since BF sings a more variable song than the WRM^62^, it is tempting to hypothesize that the changes we identified in the OT expression levels could partly influence the song nuclei, and thus differences between the BF and the WRM song.

We, lastly, hypothesize that the higher intra-strain variability in brain OT peptide content in WRM, compared to BF, could point to an evolutionary explanation according to which, out of the variable wild WRM pool, those with low brain OT peptide content were selected for breeding, to eventually make up the domesticated BF. Although such variability could also be due to differential metabolic states^63^, we believe that our experimental set up minimized the probability, since, before decapitation, all birds were kept in the dark, and did not eat, drink or sing in the hours before^64^.

In conclusion, our study revealed specific nucleotide changes in the 5’-UTR and 3’-UTR *OT* mRNA regulatory regions of the transcript within the BF in comparison with the WRM. In addition, we found a significantly lower amount of *OT* mRNA in the diencephalon of the BF compared to WRM. Whether these differences cause increased social behavior, greater song diversity, and/or a diminished stress response in the BF requires further investigation.

## Acknowledgements

We would like to thank all members of Prof. Chen, Tang and Yao lab groups for their help and support during our stay in Taiwan. We also would like to thank the field workers for their help with mist-netting birds. The present study was supported by Grants-in-Aid for Private University research Branding project and Scientific Research (JP16K18585) from the Ministry of Education, Culture, Sports, Science and Technology Japan (Y.T) and Scientific Research Grants from the Moritani Scholarship Foundation (Y.T), in Japan. CT recognised funding from the Rockefeller University; Erich D. Jarvis from the Howard Hughes Medical Institute and the Rockefeller University. CB acknowledges funding from the Spanish Science Ministry (grant PID2019-107042GB-I00).

## Data availability

The authors confirm that all of the data underlying the reported findings are included in the article. All raw data that support the findings of this study are available in the Supplementary Tables and in our publicly available Github repository (https://github.com/constantinatheo/OT-domestication).

