## Supplementary Figures for "Decreased synthesis and variable gene transcripts of oxytocin in a domesticated avian species"

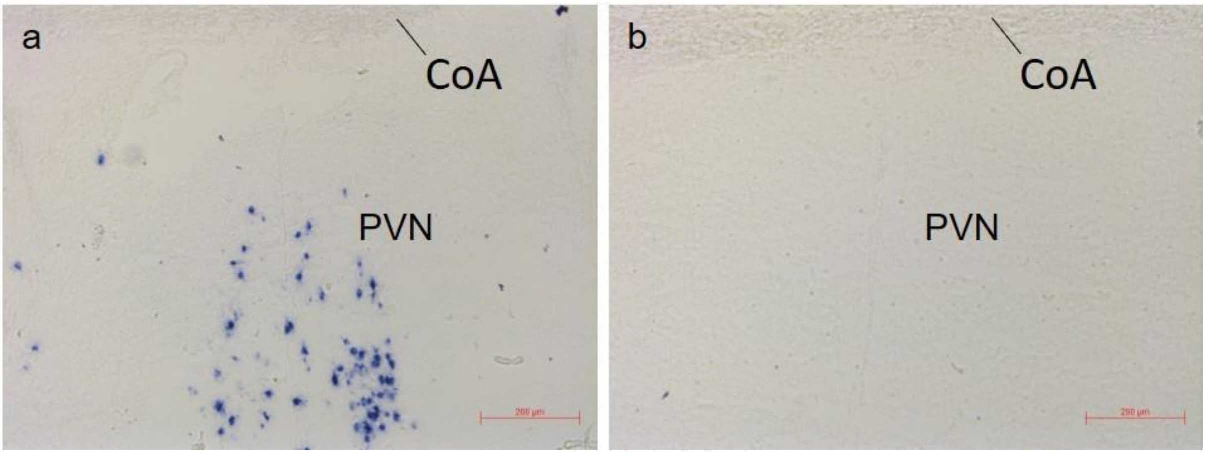


**Supplementary Figure 1: Representative sections of prepro-oxytocin (OT) mRNA expression as detected by *in situ* hybridization.** (a) Hybridization with antisense prepro-OT probe demonstrates OT mRNA expression in the PVN of Bengalese finch. (b**)** Hybridization with sense prepro-OT probe serves as negative control for the specificity of the antisense probe. CoA, anterior comissure; PVN, praventricular nucleus of the hypothalamus. Scale bars represent 200 µm.

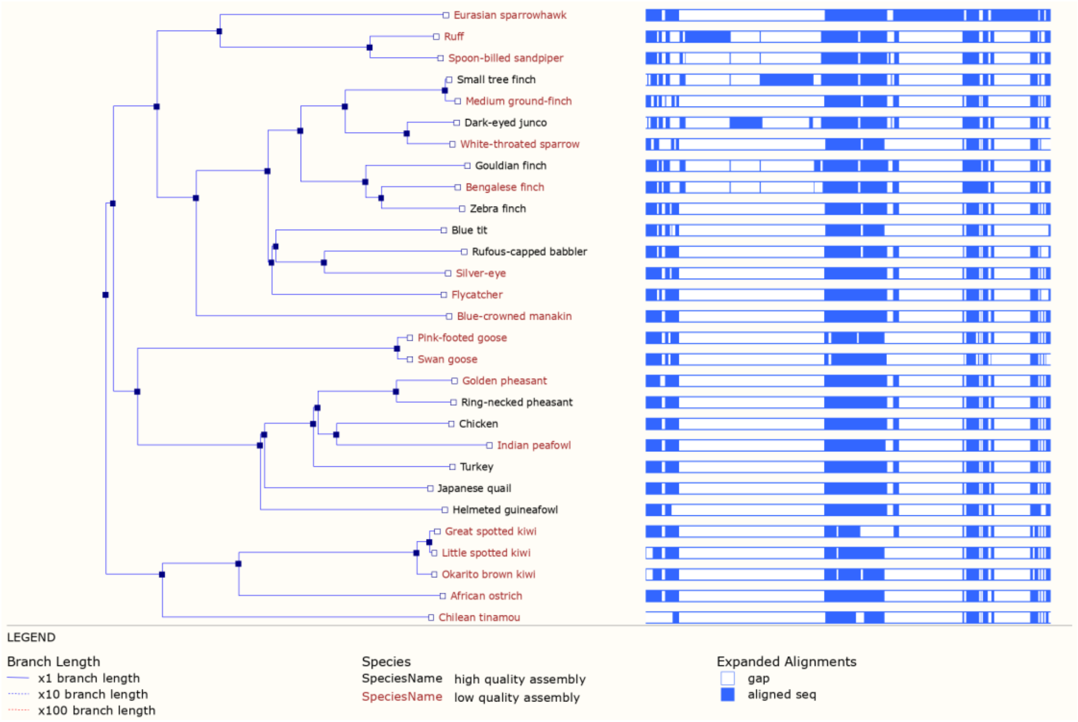


**Supplementary Figure 2:** ***OT* amino acid phylogeny**. *OT* tree topology inferred with the phylogenetic TreeFam method on an amino acid alignment generated via the Ensembl 'Gene tree' tool. Left: blue boxes, inferred speciation events. Right: Blue bars, multiple amino acid alignment made with MUSCLE; white areas, gaps in the alignment.

a

**Position:** chr4:88,651,341-88,651,341

**Total Bases in view:** 1

**Statistics on:** 1 **bases** (% 100.0000 coverage)

| Database: galGal6 | | | | | | Table: phyloP77way | | | | | |
|---|---|---|---|---|---|---|---|---|---|---|---|
| Chrom | Data start | Data end | # of Data values | Each data value spans # bases | Bases covered | Minimum | Maximum | Range | Mean | Variance | Standard deviation |
| chr4 | 88651341 | 88651341 | 1 | 1 | 1 (100.00%) | 1.02034 | 1.02034 | 0 | 1.02034 | 0 | 0 |

| 20 bin histogram on 1 values (zero count bins not shown) | | | | | | | |
|---|---|---|---|---|---|---|---|
| bin | range in bin minimum | maximum | count | Relative Frequency | log2(Frequency) | Cumulative Relative Frequency (CRF) | 1.0 - CRF |
| 15 | 0.649 | 2.0256 | 1 | 1 | 0 | 1 | 0 |

b

**Position:** chr4:88,651,382-88,651,382

**Total Bases in view:** 1

**Statistics on:** 1 **bases** (% 100.0000 coverage)

| Database: galGal6 | | | | | | Table: phyloP77way | | | | | |
|---|---|---|---|---|---|---|---|---|---|---|---|
| Chrom | Data start | Data end | # of Data values | Each data value spans # bases | Bases covered | Minimum | Maximum | Range | Mean | Variance | Standard deviation |
| chr4 | 88651382 | 88651382 | 1 | 1 | 1 (100.00%) | -0.770591 | -0.770591 | 0 | -0.770591 | 0 | 0 |

| 20 bin histogram on 1 values (zero count bins not shown) | | | | | | | |
|---|---|---|---|---|---|---|---|
| bin | range in bin minimum | maximum | count | Relative Frequency | log2(Frequency) | Cumulative Relative Frequency (CRF) | 1.0 - CRF |
| 13 | -2.1042 | -0.7276 | 1 | 1 | 0 | 1 | 0 |

**Supplementary Figure 3: PhyloP (phyloP77way) results.** PhyloP77way (UCSC Genome Browser) uses alignments (multiz) of 77 vertebrate species: 55 birds, 10 reptiles (alligator, snake, frog) and 12 other species (fish, human, mouse, lamprey) and measures evolutionary conservation using two methods (phastCons and phyloP) from the PHAST package, for all 77 species. Sites predicted to be conserved are assigned positive scores, while sites predicted to be fast-evolving are assigned negative scores. The absolute values of the scores represent -log p-values under a null hypothesis of neutral. **a,** Mean positive phylop77way score for the A > G transition site, aligned with chicken (*Gallus gallus*). Coordinates are shown for GRCg6a/galGal6. **b,** Mean negative phylop77way score for the C > T transition site, aligned with chicken (Gallus gallus). Coordinates are shown for GRCg6a/galGal6.

### Supplementary Data and Tables

Supplementary Data 1: LonStrDom1 and lonStrDom2 OT sequences.

Supplementary Data 2: LonStrDom1 vs. LonStrDom2 OT alignment.

Supplementary Data 3: OT cDNA sequences isolated from the brain of each birds

Supplementary Table 1: The origin and history of the birds used in this study.

Supplementary Table 2: Primers used for 3'-RACE, 5'-RACE, ORF cloning and in situ hybridization probe template.

Supplementary Table 3:Genome IDs and specific location coordinates of the OT gene sequences used in the multialignment.

Supplementary Table 4: Primers used for OT, OTR and PPIA in qPCR experiments.

Supplementary Table 5: Number of cells expressing oxytocin mRNA in each brain nucleus. Standard error of mean (SEM), Coefficient variation (%, CV)

Supplementary Table 6: RT-qPCR raw data used to compare relative OT mRNA expression between the BF and WRM.

Supplementary Table 7: RT-qPCR raw data used to compare relative OTR mRNA expression between the BF and WRM.

Supplementary Table 8: EIA raw data used to compare brain OT peptide content between the BF and WRM.
