## Supplementary Data S1 for "Decreased synthesis and variable gene transcripts of oxytocin in a domesticated avian species"

Ensembl ID: [ENSLSDG00000001590.1](https://www.ensembl.org/Lonchura_striata_domestica/Gene/Summary?db=core;g=ENSLSDG00000001590;r=MUZQ01000027.1:5804364-5808766;t=ENSLSDT00000002331)

>MUZQ01000027.1 dna:primary_assembly primary_assembly:LonStrDom1:MUZQ01000027.1:5804910:5806305:-1

CGCCGCTATGTCCTGCAAGGCTCTGGCTCTCTGCCTCCTGGGGCTCCTGGCTCTCTCCTC

CGCCTGCTACATCCAGAACTGCCCCATCGGGGGCAAACGTGCCGTGCTGGACATGGACAT

CAGGAAGGTAGGTCCTGGGGGGCTCGGAGCATTCCCGACCCCAGCCTTCCCTTCTCCCAA

CCTTCTCCCCCTCTGCTCCCCGAGCAGTGCCTGCCCTGCGGTCCCCGCAACAAGGGGCGC

TGCTTCGGGCCCAACATCTGCTGCGGGGAGGAGCTGGGCTGCTACATCGGCACGTCGGAC

ACGCTGCGCTGCCAGGAGGAAAACTTCCTGCCCACCCCCTGCGAGTCGGGACGCAAAGCC

TGCGGCTCCGGAGGGAGCTGCGCCGCTCCCGGCATCTGCTGCAGCACCGGTGAGGGCGGC

ACGGCATCCCGGGGTTCCTGCCGCGGAGCCCTCGCACAGCCCGGGGGGAATTCAGGGTTC

TGAGAACAGCGGGAGAGGGGAGGATGCCCAAGGGATTGGCTCCGTCCCATCCGTGTGTCC

ATCCATCGCCCCGACAGCCAGACCACGCTCCGTGAGCTATGCAGACCCCAGGGAAGGGAT

CCTGGGAGAACAGGAGGGTTTGGGGCAGAGGGAAGCGGATAGCAAACAGATTTTGGGGGC

TGAGCAAGTGCAGGAGCCACATTCCTTCCTGTTCTCCGTTATTCTCAAAATCCCACCCTC

CTCCTTAAATTAAACTACCAGAGATGCTGCTATAAATCCTGGGAAGCATCCAGAGAATGG

AGTGAGCACAAGCTCATTAAACCCCAGTGCTGGGGGGAGGGTCTCTGGGGGCTGTTTCAG

GGGGTCTCTGAATGTTTGGGAGATTTCTGGGGGCAGTGGTTTTGGGGGTCCCTGGGGACT

GTTTTGGGGGTTCCCTGGGAGCTGGTTCAGAGGGTCCCTGGGGGACTGTTTCAGGGATCT

CTGGGAGCTGTTTTGGGCGTCCCTGGGGGCTGTTTTAGGGATCTCTGGGGGTTATTTTGG

GGGTTCCCTGTGGGCTAGTTCAGGGATCTCTGGGAGCTGTTTCAGGGATCCCTGGGAGCT

CTTTTGGGTGTTCCTGGGGGCTGTTTTGGGGGTTCTCTGGGAGCTGGTTCAGAGGGTCCC

TGGGGCCTGTTTCAGGGATCTCTGAGGCCCATTTTGGGGGGGTCCCTGGGGTGCTGTCTG

GTTTCACACCACACTTCCCCCTCTCTCCCCTCAGAGGGCTGTGGCACTGACTCATCCTGT

GACCAGGAGATGCTGTTTGTGTAGCCCACCCCGGAGAGAATCCGCAGGATCCCGCTTCCA

TCGCTGTCCCAGCCCTGGGCTGTGACTCAGACTGAAGTGATGAGTTAATTAGAAATAAAA

CTTGGACAGAAAAACA

GCT / MUZQ01000027.1: 5,804,914-5,804,916

G A +1303 MUZQ01000027.1:5,806,213

T C +1335 MUZQ01000027.1:5,806,245

>NC_042569.1:68957139-68958557 Lonchura striata domestica isolate Mets1 chromosome 4, lonStrDom2, whole genome shotgun sequence

CCTGGGCTGTGACTCAGACTGAAGTGATGAGTTAATTAGAAATAAAACTTGGACAGAAAAACAACCCGGACAGGAGATGCTGTTTGTGTAGCCCACCCCGGAGAGAATCCGCAGGATCCCGCTTCCATCGCTGTCCCAGCCTGTCTGGTTTCACACCACACTTCCCCCTCTCTCCCCTCAGAGGGCTGTGGCACTGACTCATCCTGTGACCTGGTTCAGAGGGTCCCTGGGGCCTGTTTCAGGGATCTCTGAGGCCCATTTTGGGGGGGTCCCTGGGGTGGAGCTGTTTCAGGGATCCCTGGGAGCTCTTTTGGGTGTTCCTGGGGGCTGTTTTGGGGGTTCTCTGGGAGTGGGGGCTGTTTTAGGGATCTCTGGGGGTTATTTTGGGGGTTCCCTGTGGGCTAGTTCAGGGATCTCTGGCCTGGGAGCTGGTTCAGAGGGTCCCTGGGGGACTGTTTCAGGGATCTCTGGGAGCTGTTTTGGGCGTCCCGTCTCTGAATGTTTGGGAGATTTCTGGGGGCAGTGGTTTTGGGGGTCCCTGGGGACTGTTTTGGGGGTTCAGAATGGAGTGAGCACAAGCTCATTAAACCCCAGTGCTGGGGGGAGGGTCTCTGGGGGCTGTTTCAGGGGCTCAAAATCCCACCCTCCTCCTTAAATTAAACTACCAGAGATGCTGCTATAAATCCTGGGAAGCATCCAGAGCGGATAGCAAACAGATTTTGGGGGCTGAGCAAGTGCAGGAGCCACATTCCTTCCTGTTCTCCGTTATTCACGCTCCGTGAGCTATGCAGACCCCAGGGAAGGGATCCTGGGAGAACAGGAGGGTTTGGGGCAGAGGGAAGAGGGGAGGATGCCCAAGGGATTGGCTCCGTCCCATCCGTGTGTCCATCCATCGCCCCGACAGCCAGACGCATCCCGGGGTTCCTGCCGCGGAGCCCTCGCACAGCCCGGGGGGAATTCAGGGTTCTGAGAACAGCGGGCAAAGCCTGCGGCTCCGGAGGGAGCTGCGCCGCTCCCGGCATCTGCTGCAGCACCGGTGAGGGCGGCACGACATCGGCACGTCGGACACGCTGCGCTGCCAGGAGGAAAACTTCCTGCCCACCCCCTGCGAGTCGGGACGCCCTGCGGTCCCCGCAACAAGGGGCGCTGCTTCGGGCCCAACATCTGCTGCGGGGAGGAGCTGGGCTGCTTCGGAGCATTCCCGACCCCAGCCTTCCCTTCTCCCAACCTTCTCCCCCTCTGCTCCCCGAGCAGTGCCTGCAGAACTGCCCCATCGGGGGCAAACGTGCCGTGCTGGACATGGACATCAGGAAGGTAGGTCCTGGGGGGCCGCTATGTCCTGCAAGGCTCTGGCTCTCTGCCTCCTGGGGCTCCTGGCTCTCTCCTCCGCCTGCTACATCAAATCTCCCCCTGTCCCGC

>4 dna:chromosome chromosome:GRCg6a:4:88649940:88651459:-1

AAGCATCCTGATCCTGCAGCCATGTTCTACAAGGCGCTCACCGTCTGCCTGCTGGGGCTT

CTGGCTCTCTCCTCAGCTTGTTATATCCAGAACTGCCCCATCGGGGGTAAGCGTGCCGTG

CCAGACATGGACATCAGGAAGGTAGGTCCCTGTGGGCTGGGTGGGCTGTGAGAGCACCTG

TCTTCCTTCTGCCACCCTTCCTTGGGCAGCTCTGAGCCTCCCGAACAGCAAGGCAAGGTG

CATGGAGCAGGGATGTTATAGGAATGGGGTTTCAGAATGGGGAATCTCAAAGAAAAGCGG

TTTGGGGATGGGCAGCCAAAAGGGAGCTGCAAAGAAGGCAGCCTGAAGCAGGATGCTGAT

GAAAACCCATCTCTCTGACAATTGCTTTTTCCCAACACTGGGAAAAAACTACGATTCCAA

TGGGGCGGTGGCACCATTTTTATGTAGAATGTTAACTTCTGGTGGGTGAATTTGGGGCAT

GATCAGCCATTTGACTAAAGCCAGGACTGGCGTGGAATCATTCAGGTTGGAAAAGACCTC

TAAGAGCACCAAGTCCAACCCCAGCACATCCCTCAGTGCCACATCTCCAGTTCTGCAACA

CCTCCAGGTTTGGTGATTTCACCACTCCTTGTGCTGCCTGTGTCAGTGCCTGACACCTCT

TTTGGGGAATTTTTTCCTAATATCTAACCTGACCCTCCACCGGTGCAGTGTGAGGCCATT

CCTTCTTATCCTACCACCGATACCTGGCAGAAAAGACCAATGCCTTCCCACCACCCCTCA

AAATATACATGAAAACAAACCCAGAGCCTTCCAGAGCAAACCACCGGGGTGAGGACAGAG

CTGAGAGCAAGATGAGGATACACACACACCGGCAGAGGGCAAAGAGATCAGGCTCAATCC

TTAACACCGTTCCTCAATTCCCTGAGCAGTGCCTGCCCTGCGGCCCCAGGAACAAGGGCC

ACTGCTTTGGCCCCAACATCTGCTGCGGGGAGGAGCTGGGCTGCTACCTGGGCACCTCTG

AAACCCTGCGGTGCCAGGAGGAGAACTTCCTGCCAACGCCCTGTGAGTCTGGGAGGAAAG

CCTGCGGCGAGGATGGGGCGAGCTGTGCAGCACCAGGAATCTGCTGCAGCAGTGGTGAGT

AACGTGGGCAGATTGGGGGCCTCACATCCACCCCAGGGCTATGGTACAGATGAGGGAGGG

GAGGATGTTACTGAGGAGGGACATGTTGCTGCTGCAGGGCACCGAGATAATGCCATGTTG

TTTGATGTTATTAAAAACAGCACAGGGAGGGGGCTCTGAGGGCTGTTTTGTGCTGATTTC

CTCTTCTCTCCCTGAGCAGAGGGCTGTGTGCTTGACTCATCTTGCAACCAAGAAATGCTG

TTTGCATAGAGGGAAAAACCAGCAGGATCTGCCATCCCATGGGCTGCACCAGCGCAGCAC

CCAGCACCATGACTCAGACTGAAGTGATGCTCCTAATTAGAAATAAAACTTGGACAGAAT

TAAAAAAAGTGTGTTTTTCT

G 88649940+1401 – 88651341

A (T) 88649940+1442---88651382---3
